# The Neurocognitive efficiency score: Derivation, validation, and application of a novel combination of concurrent electrophysiological and behavioral data

**DOI:** 10.1101/2025.01.10.632377

**Authors:** Michael J. Wenger, James T. Townsend, Sarah F. Newbolds

## Abstract

Perceptual and cognitive work require the expenditure of brain energy, and the efficiency of that energy expenditure can vary as a function of a range of exogenous and endogenous factors. The same is true for physical work, and tools have been developed to quantify energetic efficiency in the performance of physical work. The same is not true for the study of perceptual and cognitive work. We here draw on two lines of research—the formal characterization of workload capacity in perception and cognition, and the study of physical work—to propose a novel measure of neurocognitive efficiency. This neurocognitive efficiency score (NES) combines reaction time data with measures on concurrently acquired electroencephalographic (EEG) data to form a ratio that can be interpreted in terms of work accomplished relative to energy. We consider three measures on the EEG and show that one of them—global field power (GFP)—evidences a strong relationship to measures of metabolic energy expended. We then show how the NES can provide insights into differences in perceptual performance as a function of biological state. We argue that the NES has the potential for a wide range of applications in the study of perception and cognition in the context of factors including (but not limited to) aging, disease, stress, and differences in levels of expertise.

## Introduction

Brain activity in support of perception and cognition—maintenance of the resting membrane potential, generation of action potentials, and the integration of excitatory and inhibitory inputs in synaptic transmission—comes with a metabolic cost (Özugur, Kunz, & Straka, 2020). Specifically, the brain requires significant amounts of oxygen in order to generate adenosine triphosphase (ATP) in the mitochondria (e.g. Ames, 2000; Hall, Klein-Flügge, Howarth, & Attwell, 2012). A natural question that arises from these facts has to do with the nature of the relationship of that oxygen consumption to the amount of perceptual and cognitive work that can be accomplished: a question of efficiency. This is because it cannot be a priori assumed that efficiency is optimal in all cases. Obvious examples include variations in efficiency as a function of biological status (e.g., disease, age, etc.), behavioral status (e.g., level of expertise, level of fatique, etc.), environmental status (e.g., variations in external noise levels, temperature, altitude, exogenous stress, etc.), and task-specific workload. The issue of efficiency in the expenditure of biological energy in the performance of perceptual and cognitive work has long been overlooked in cognitive science and cognitive neuroscience, as the traditional focus has been on computational questions.^1^

As is discussed below, attempts to address the efficiency of energy expenditure in the performance of perceptual and cognitive work is both highly delimited and saddled with a set of methodological challenges. We here propose a novel approach to quantifying neurocognitive energy expenditure, making use of proxy measures for metabolic energy expenditure extracted from electroencephalographic (EEG) data and probabilistic measures of work accomplished based on reaction time (RT) distributions. The work draws inspiration from two lines of research. The first is the long line of work quantifying the construct of *capacity*, roughly variations in performance (as assessed using RTs) as a function of variations in workload (e.g., Townsend & Ashby, 1978; Townsend & Nozawa, 1995). The second is the line of work quantifying the construct of *energetic efficiency* in the performance of *physical* work as a function of levels of systemic iron (Haas & Brownlie, 2001; Haas, 2006; Luna, Pompano, Lung’aho, Gahutu, & Haas, 2020).

We structure our presentation as follows. We begin with an overview of the formal work done to characterize the construct of capacity in perceptual and cognitive work and provides an instantaneous measure of work being accomplished. This establishes the motivation for the first component (the numerator) of our proposed measure of neurocognitive efficiency. We then provide an overview of the way in which efficiency in the expenditure of biological energy has been quantified in the specific context of iron deficiency, the most-prevalent micronutrient deficiency in the world (*Anemia*, 2024). We next consider a set of candidate measures for metabolic energy expenditure during the performance of perceptual and cognitive work that can be extracted from EEG data. Using an analysis of previously unpublished data from an earlier study (Wenger, DellaValle, Murray-Kolb, & Haas, 2017), we show that one of those candidate measures—global field power (GFP, Skrandies, 1989, 1990)—is directly related to measures of metabolic energy expended. This establishes the motivation for the second component (the denominator) of our proposed measure of neurocognitive efficiency: the ratio of work accomplished to energy expended. We close by showing how this novel neurocognitive measure provides an important insight into differences in efficiency in perceptual and cognitive work as a function of biological status, specifically systemic iron status, in application to previously unpublished data from an earlier study (Newbolds & Wenger, 2024).

### Characterizing capacity

The construct of capacity has a long history in scientific psychology, referring in various contexts to hypothesized limits on storage (e.g., Cunningham, Yassa, & Egeth, 2015; Fukuda & Vogel, 2019), speed of processing (e.g., Kail & Salthouse, 1994), processing resources (e.g., Fisher, 1984; Wickens, 1984), and combinations of ideas regarding storage and processing (e.g., Cowan, 2012; Oberauer, Farrell, Jarrold, & Lewandowsky, 2016), among others. The particular perspective that we adopt here draws on the mathematical characterization of capacity with respect to changes in performance (specifically reaction times, RTs) as a function of variations in perceptual and cognitive workload, referred to as workload capacity (e.g., Blaha & Houpt, 2015; Heathcote et al., 2015; Houpt & Townsend, 2012; Townsend & Eidels, 2011; Yu, Chang, & Yang, 2014). While this characterization of capacity was given its most comprehensive treatment in the development of systems factorial technology (Townsend & Nozawa, 1995), a critical initial proposal for characterizing capacity for accomplishing work was provided by Townsend and Ashby (1978, pp. 217 ff), who suggested the potential of using the hazard function on the RT distribution (see also Townsend & Ashby, 1983, pp. 248 ff). The hazard function is also known as the intensity function, or the age-specific failure rate (Cox, 1962; Finkelstein, 2008; McGill & Gibbon, 1965).

The formal definition of the hazard function is as a limit on a conditional probability as a function of time

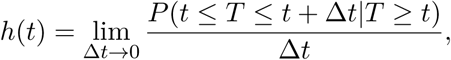

which is also sometimes expressed as a ratio

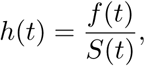

in which the numerator is the probability density function and denominator is the survivor function, which itself is the complement of the cumulative distribution function. The hazard function quantifies the probability that some event will happen in the next instant of time, given that it has not happened yet. In the context of a perceptual or cognitive task in which RTs are measured, this would be the probability of making a response in the next instant of time given that response has not yet been made. If we interpret the time required to make a response as an indicator that the requisite perceptual or cognitive work has been completed (including the time to encode the stimulus, generate a motor signal, and execute the observable response), then the allowable interpretation of *h*(*t*) is as an instantaneous measure of the amount of work that has been accomplished. Building on this interpretation, it is then possible to consider the integral of the hazard function

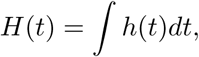

which is estimable from data as

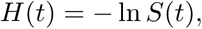

where *S*(*t*) is the survivor function of the RT distribution. The integrated hazard function can then be interpreted as the *cumulative* amount of work accomplished up to some time *t*. This function *H*(*t*) has formed the basis for quantitative characterization of workload capacity in systems factorial technology (Little, Altieri, Yang, & Fific, 2017; Townsend & Nozawa, 1995; Townsend & Wenger, 2004; Townsend & Eidels, 2011; Townsend & Altieri, 2012).

In developing systems factorial technology, Townsend and Nozawa (1995) showed that it was possible to strongly adjudicate the long-standing issue of serial vs. parallel processing given two or more inputs, but that doing so required consideration of four fundamental characteristics of information processing:^2^ architecture, stopping rule, independence in rate, and capacity. Figures 1 and 2 provide schematic representations of these four characteristics.

**Figure 1.**
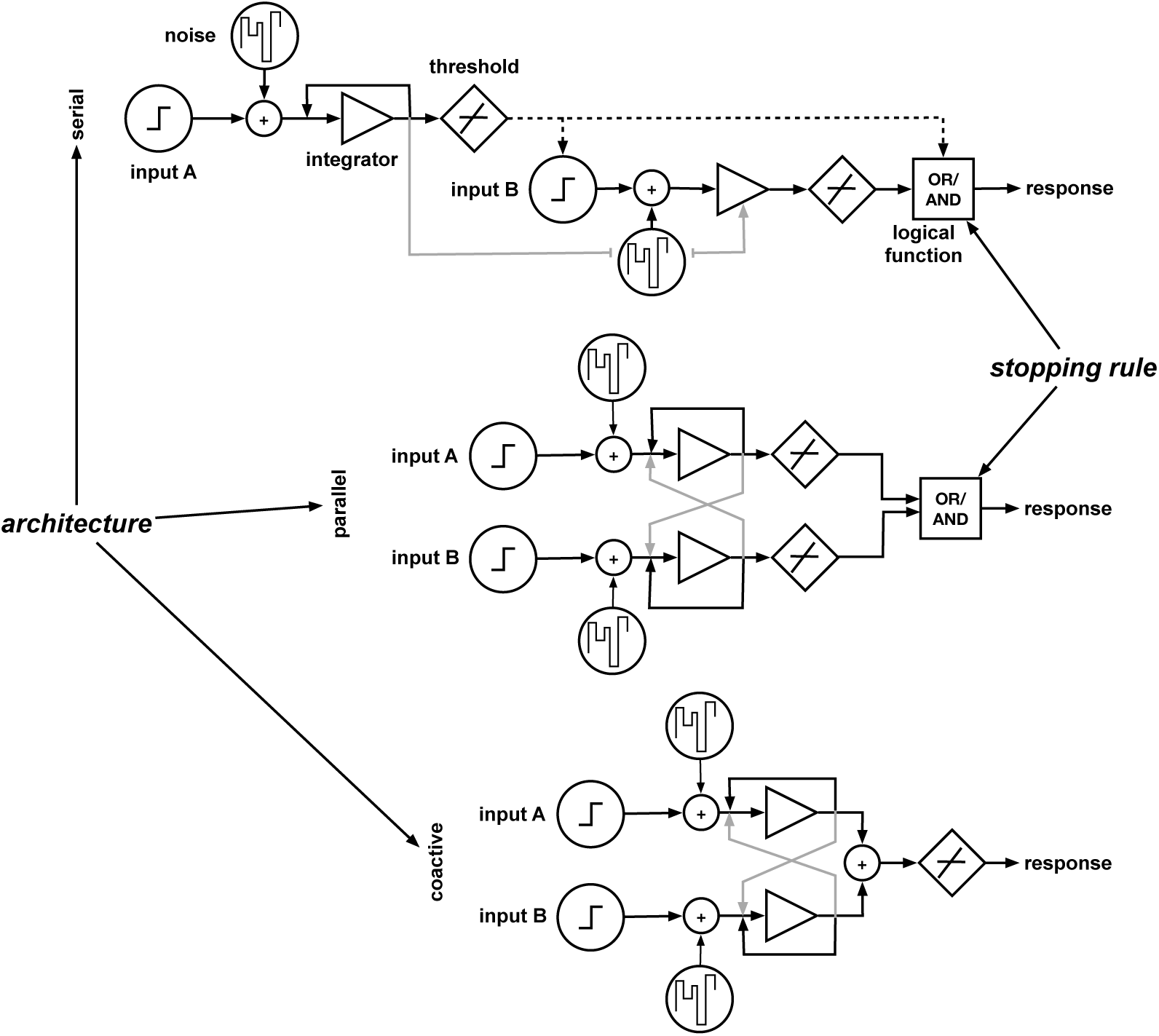
Schematic representation of three of the four processing characteristics identified by Townsend and Nozawa (1995) being necessary to consider in characterizing an information processing system operating on two or more inputs: architecture, stopping rule, and independence/dependence in rate of processing. Note that the light gray lines indicate possible sources of dependency in rate of processing.

**Figure 2.**
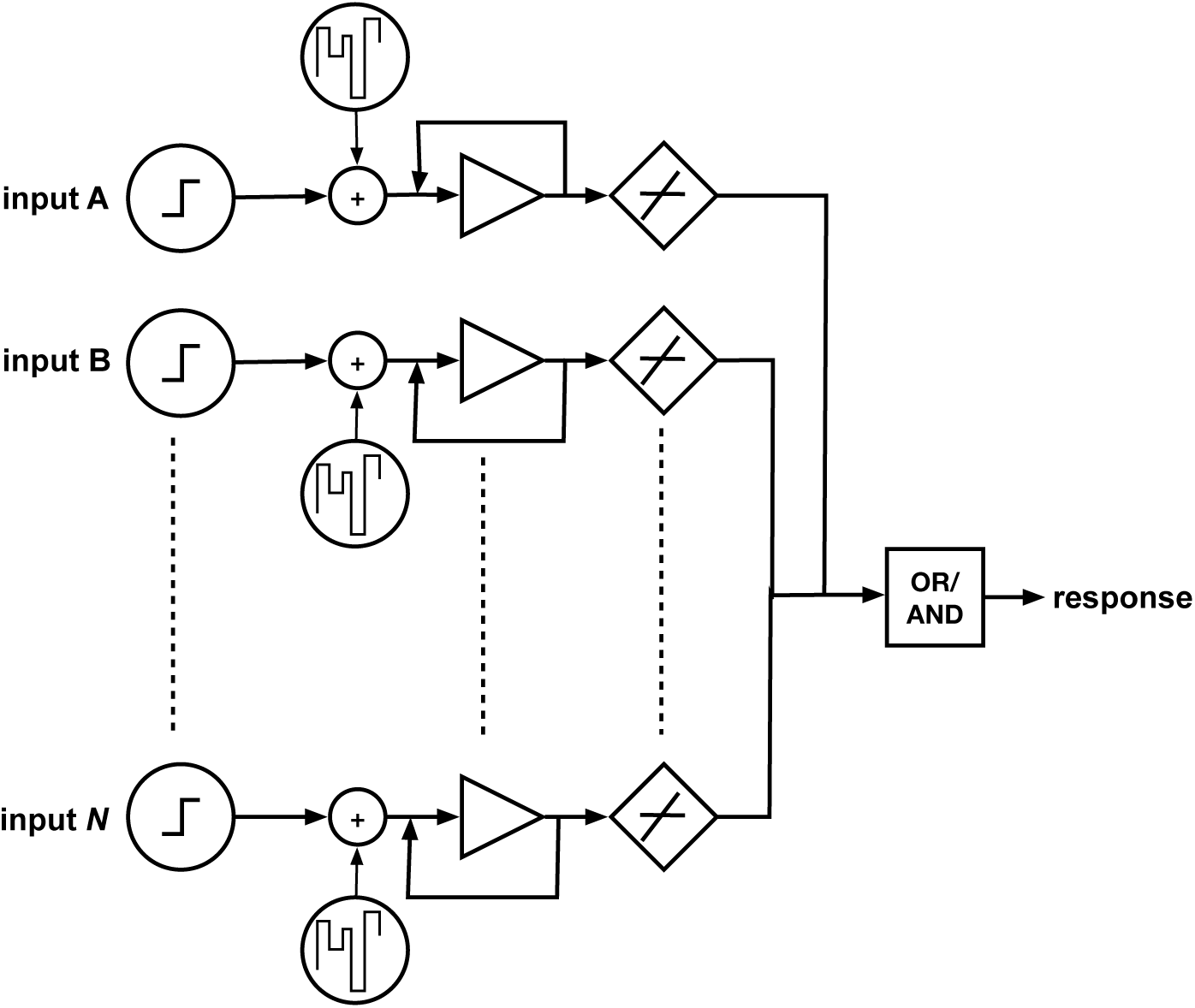
Schematic representation of the construct of workload capacity and the basis of the capacity coefficient as developed by Townsend and Nozawa (1995). Note that independence in rate of processing is assumed.

With respect to architecture, Townsend and Nozawa (1995) identified three possibilities: serial, parallel, and a variant of a parallel architecture known as coactivation, in which information in the parallel channels is pooled prior to comparison to a threshold. Stopping rule refers to the amount of processing that needs to be completed prior to generating a response, with the two alternatives being self-terminating (corresponding to the Boolean OR operator) and exhaustive (corresponding to the AND operator). What also must be considered is the extent to which the rates processing in each of the channels are either independent or dependent.

Of critical concern for the present discussion is the fourth characteristic, workload capacity, with the characterization being framed with respect to changes in performance with changes in workload. This is represented schematically in Figure 2 by additional inputs. Townsend and Nozawa (1995) noted that if it is assumed that the system in question is parallel, with independent channels, and an OR stopping rule, then a critical regularity holds: the integrated hazard function for the time required to process all the inputs is identical to the sum of the integrated hazard functions for the time required to process each of the individual inputs (see also Townsend & Ashby, 1983, pp. 248 ff). This the basis for the capacity coefficient:

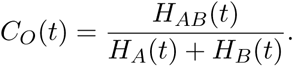

Here the O in the subscript indicates that this is specific to systems with an OR response rule and, with the assumptions just stated, *C_O_*(*t*) will be equal to 1. In the interpretation developed earlier, this means that the cumulative amount of work accomplished given two inputs is the same as the sum of the cumulative measures of work for the two inputs separately, and quite naturally indicates unlimited capacity processing. Values *<* 1 indicate that as workload is increased, performance degrades, indicating limited capacity processing, while values *>* 1 indicate that performance improves as workload is increased, indicating super-capacity processing. Note that while the capacity coefficient is framed here for two inputs, the formalism can be generalized to *N* inputs. Also note that a corresponding capacity coefficient for a system with parallel independent channels with an AND stopping rule was developed by Townsend and Wenger (2004). The utility and interpretability of *H*(*t*) provides an important hint as to what might be useful and interpretable in developing our measure of neurocognitive efficiency.

Before moving on, we should note that hazard functions, or variants, have been used in at least two additional ways of characterizing information processing systems. Lappin and colleagues (Lappin, Morse, & Seiffert, 2016; Lappin, Seiffert, & Bell, 2020; Lappin & Bell, 2021) have suggested that, rather than estimating the cumulative hazard function (and then its derivative) by way of the natural log, estimating it using log_2_, which allows for interpretation in terms of bits per sec. They have used this approach to demonstrate an invariance across stimulus conditions that appears to be consistent with Shannon’s (1948; 1963) theorem that that any physical channel has a maximum rate at which it can transmit information. An additional perspective is provided in the work of Chechile (2003, 2011, 2022), which demonstrates that the reverse hazard function (Woodroofe, 1985) can be effectively used in combination with the hazard function to distinguish alternative models of processing at the level of the distribution of processing times.

### Characterizing energetic efficiency in the performance of physical work

A model for how we can develop a measure of efficiency for perceptual and cognitive work comes from resaerch that has quantfied energetic efficiency in the performance of physical work. Specifically, Haas and colleagues have shown that there is regular and reliable relationship between energetic efficiency and levels of systemic iron in women of reproductive age (e.g., DellaValle & Haas, 2014; Haas & Brownlie, 2001; R. Li et al., 1994; Y. I. Zhu & Haas, 1997; Y. Zhu & Haas, 1998), with lower levels associated with lower energetic efficiency and with improvements in energetic efficiency being possible with iron repletion (Y. Zhu & Haas, 1998).

To illustrate how this is quantified, we use results from a study by Luna et al. (2020). In brief, that study was conducted in Rwanda and examined the potential impact of consuming an iron-biofortified^3^ bean, a staple food in Rwanda, on systemic iron levels in college-aged women in Rwanda who were experiencing iron deficiency either with or without anemia.^4^ A subset of the women participating in that study performed a physical performance task on a cycle ergometer following a standard protocol in exercise science (Cooper & Storer, 2001) at the beginning (baseline) and end (endline) of an 18-week period during which they consumed either the biofortified bean or a comparison bean. Physical workload in the task was manipulated by increasing the resistance on the cycle ergometer. Women performed this task while being fitted with a portable metabolic measurement system which assessed the amount of oxygen consumed and the amount of carbon dioxide expired (among other quantities). Using these values, it is possible calculate energy expended (EE) as

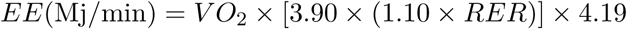

(de V. Weir, 1949). Work efficiency (WE) can then be calculated as

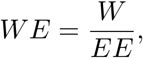

a ratio which characterizes relationship between the amount of work performed (in this case, in Watts) and the amount of energy expended.

In brief, the results from Luna et al. (2020) were as follows. At baseline, the women in the two groups (biofortified vs. comparison bean) showed equivalent levels of work efficiency. At endline, women who had consumed the biofortified bean had higher levels of both hemoglobin (Hb, the measure of iron related to oxygen transport) and serum ferritin (sFt, the storage form of iron, also related to synthesis and regulation of neurotransmitters, including dopamine, serotonin, and glutamate) than did the women who consumed the comparison bean. Critically for present purposes, when considering all of the women who consumed the biofortified bean, increases in Hb were reliably associated with increases in work efficiency at the highest level of workload, a relationship which was also observed for the women who were both iron deficient and anemic at baseline. For the women who were iron deficient but not anemic at baseline, increases in work efficiency at the highest level of workload were reliably associated with increases in sFt.

The ability of this approach to quantify work efficiency and improvements in efficiency with iron supplementation provides a model for how we might quantify neurocognitive efficiency. In order to do this, we need variables that correspond to those in the numerator and denominator of the equation for work efficiency. As suggested above, the hazard function and its integral provide measures that can be interpreted in terms of work performed, either instantaneously or cumulatively. Which raises the question of how we might quantify brain energy expenditure.

### Prior work characterizing brain energy expenditure

Previous attempts at indirectly estimating brain energy expenditures as a function of mental workload have had a number of limitations. First, indirect measures such as cerebral blood flow and blood oxygen concentration are limited in the range of covariation with cortical activity. This appears to in part be due to the fact that the brain, even at rest, is quite active metabolically, such that most increases in task demands result in small increases in overall oxygen requirements and energy demands (Lin, Fox, Hardies, Duong, & Gao, 2010; Raichle & Gusnard, 2002; Sokoloff, 2004). In addition, task-dependent changes in cerebral blood flow seem to reflect a generalized mobilization of brain resources rather than a task-specific increase, with some studies suggesting that the differential between cerebral blood flow and cerebral oxygen metabolism is roughly an order of magnitude, if not more (Fox & Raichle, 1986; Fox, Raichle, Mintun, & Dence, 1988; Raichle & Gusnard, 2002). Second, global measures of brain energy demands are too coarse, spatially and temporally, to reflect the short-duration engagement of specific brain regions in processing that may be complete within hundreds of ms (Lin et al., 2010; Raichle & Gusnard, 2002; Sokoloff, 2004). Third, almost all published efforts lack a systematic analysis, specification, or modeling of the specific brain regions that are hypothesized to support task performance. Taken in combination with the coarseness of the global measures of brain energy expenditure, this lack of specification means that there is no guidance as to where to measure and limited information in what can be measured. Fourth, the range of workload manipulations that has been used is extremely limited, sometimes being restricted to comparisons involving the difference between the presence and absence of work (e.g., Garde, Laursen, Jørgensen, & Jensen, 2002; Petrek, 2008, 2009). Such restrictions limit the ability to quantify changes in performance as a function of changes in level of demand, and run the risk of considering levels of demand that may may not be sufficiently taxing, a problem that plagued early attempts to behaviorally quantify capacity effects in perception and cognition (as discussed in, e.g., Fisher, 1982, 1984). Finally, the standard behavioral measures of performance—-such as mean accuracy and latency—-fail to provide the necessary variation within individual subjects needed to allow quantification of individual responses to changes in external workload. Finally, it should be noted that other measures of brain energy expenditure, such as high field magnetic resonance imaging or functional near-infrared spectroscopy (e.g., Hyder, 2004; Hyder & Blumenfeld, 2004; Laughlin & Attwell, 2004; Lin et al., 2010; Raichle & Gusnard, 2002; Sokoloff, 2004) can potentially address some of these problems, but also have limitations, including lack of temporal resolution and potential loss of temporal information. The potential for using EEG as a basis for the type of indirect calorimetry that we are envisioning comes from the fact that glucose utilization in the brain is directly related to the intensity of neural activation, in particular the generation and propagation of action and generator potentials and the release, uptake, and re-uptake of neurotransmitters (e.g., Fairclough & Houston, 2004; Fox et al., 1988; Lin et al., 2010; Raichle & Gusnard, 2002; Sokoloff, 2004). Activation of neural tissue produces increases in oxygen consumption and glucose metabolism that are nearly linearly related to spiking frequency (e.g., Hyder, Rothman, & Shulman, 2002; Laughlin & Attwell, 2004; Özugur et al., 2020). Given that activity at the synapses—the apical dendrites in particular—and overall spike frequency are among the determinants of many features of EEG and event-related potentials (e.g., amplitudes of component features in the time domain and spectral power in the frequency domain, see Nunez & Srinivasan, 2006), along with the ability to measure EEG at time-scales that are commensurate with the durations over which perceptual and cognitive processing occurs, suggests that EEG may be potentially sensitive to task-dependent onset and duration of neural activity. In addition, there are precedents for using aspects of EEG data as metrics for effort and fatigue during cognitive work (Berka et al., 2007; Dasari, Shou, & Ding, 2017; Jung & Lee, 2007; Longo, Wickens, Hancock, & Hancock, 2022).

### Proposal for the neurocognitive efficiency score

As a reminder, our goal is to obtain a measure of neurocognitive energetic efficiency that can be used in the study of perceptual and cognitive performance that has the same form as the calculation of work efficiency. As noted above, a natural choice for the numerator is either *h*(*t*) or *H*(*t*), given the interpretation in terms of work accomplished, either instantaneously or cumulatively. Given that we propose to use measures on EEG as proxies for brain metabolic energy, and given the temporal resolution of EEG, we opt for *h*(*t*) as an instantaneous measure of work accomplished.

Three possibilities can be considered for the denominator, one in the time domain and two in the time-frequency domain.^5^ In the time domain, we consider the global field power (GFP) of the EEG, calculated as the spatial standard deviation of the mean across all electrodes at each time point, with the units being microvolts (*µ*V). The GFP was championed for use with EEG by Lehmann and Skrandies (1980; 1982; 1989, 1990) and has been used to identify time ranges for the extraction of peaks in ERPs (e.g., Halpern, Martin, & Reed, 2008; Centanni, Halpern, Seisler, & Wenger, 2020; Von Der Heide, Wenger, Bittner, & Fitousi, 2018). Skrandies suggested that the GFP could be interpreted as indicator of synchronous neural activity, with the peak of the GFP indicating “the occurrence time of maximal activity in the potential distributions, reflecting the synchronous activation of a maximal number of neurones” (Skrandies, 1990, p. 138). This interpretation of the GFP suggests its possible utility as an indirect indicator of brain energy expenditure.

The two possibilities for the denominator in the time-frequency domain are summed positively rectified power in the *α* (8-13 Hz) and *γ* (30-90 Hz) bands, given the relationships that have been noted for activity in these bands and different aspects of attention (Borghetti, Morris, Jack Rhodes, Haubert, & Veksler, 2021; Doesburg, Roggeveen, Kitajo, & Ward, 2008; Landau, Esterman, Robertson, Bentin, & Prinzmetal, 2007; Mizokuchi, Tanaka, Sato, & Shiraki, 2024; Noah et al., 2020; Souza & Naves, 2021). When the units of time-frequency analyses are decibels (dB), values less than 0 indicate reductions in activity while values greater than 0 indicate increases in activity. Given that we are interested in possible increases in energy expenditure, we positively rectify the time series and then sum the power across the range of frequencies in each band.

Our proposed measure of neurocognitive efficiency can then be formed by taking the ratio of *h*(*t*) to one of the possible measures energy expenditure, for example

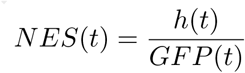

and the two additional alternatives being

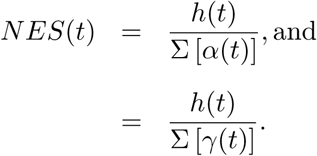

In each case, we have a measure that can be interpreted in the same way as the measure of work efficiency described above. There is a crucial step that needs to be taken before this measure can be meaningfully used. Specifically, we need to know whether any of the three possible measures in the denominator are regularly related to metabolic energy expenditure during the performance of cognitive work and, if more than one of them are, which has the strongest relationship. This is the issue to which we turn next.

### Validation of the neurocognitive efficiency score

In order to know whether or not any of the three forms of *NES*(*t*) are interpretable in the manner intended, we need to be able to examine the possible relationships between the three possibilities for the denominator and metabolic energy expenditure during the performance of cognitive work. To do this, we need three concurrent measures: RTs, for the calculation of *h*(*t*); EEG, for the calculation of *GFP* (*t*), Σ [*α*(*t*)], and Σ [*γ*(*t*)], and the volumes of oxygen consumed and carbon dioxide expired. It turns out that we do have all of these measures, collected simultaneously, in unpublished data from an earlier study of the effects of iron deficiency on cognitive performance in women of reproductive age (Wenger, DellaValle, et al., 2017).

In that study, we sought to perform a cognitive analog to the exercise task used by Luna et al. (2020). Specifically, we sought to design a cognitive task in which the initial workload was light and then systematically increased until performance degraded to chance. We had participants perform the task, during which we recorded choice frequencies and RTs, and concurrently recorded EEG, volume of oxygen consumed, and volume of carbon dioxide expired.

### Description of the experimental method

In brief, the methods were as follows (interested readers should consult Wenger, DellaValle, et al., 2017, for methodological details). Participants were women of reproductive age who were screened by way of blood tests to be in one of two groups: (a) iron deficient and not anemic (IDNA), defined as sFt *≤* 16.0 *µ*g/L and Hb *≥* 12.0 g/dL; and (b) iron sufficient (IS), defined as sFt *≥* 20.0 *µ*G/L and Hb *≥* 12.0 g/dL. Participants in the IS condition were matched to those in the IDNA condition on age, education level, and self-reported activity level.

The behavioral task was a combination of a visual Sternberg (1966) short duration memory task and a simple mental math task. We used this dual task paradigm as we discovered in pilot work that the Sternberg task was not sufficiently difficult on its own to allow performance to be degraded to chance. The Sternberg task involved the presentation of one to four “squiggle” stimuli (based on Blaha, Busey, & Townsend, 2009, see Figure 3). The difficulty of the task was varied by factorially manipulating the number of squiggles to be remembered (set size: 1, 2, 3, or 4) and the complexity of each squiggle (number of segments: 1, 2, 3, and 5). The task started at the lowest level of difficulty (set size 1 with 1 segment), with difficulty increasing in a step-wise manner, first by increasing the number of segments at each set size. Once the number of segments reached the maximum for the current set size, set size was increased to the step and the number of segments reset to the lowest level. This was repeated until the highest level of difficulty (set size 4 with 5 segments) was reached.

**Figure 3.**
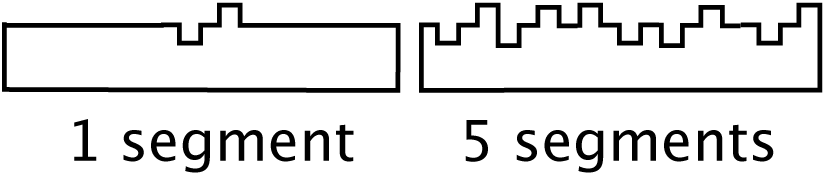
Examples of the to-be-remembered stimuli used in the visual Sternberg task in Wenger et al. (2017). Adapted from Figure 1 in Wenger et al. (2017)

Each trial in this task was self-initiated and began with a fixation cross followed by the presentation of the to-be-remembered squiggles at the current level of set size and number of squiggles. Each presentation of a squiggle was accompanied by the presentation of a single-digit number to the side of the squiggle, with participants required to mentally maintain a cumulative total of the numbers. Following the presentation of the last squiggle and digit, a test squiggle was presented that was either identical to one of the to-be-remembered (old) squiggles or different from one of the presented squiggles (new) by one segment, with participants required to indicate whether the test item was old or new. A total of 30 old and 30 new trials were presented at each combination of set size and number of segments. Immediately after the test squiggle, a test sum was presented that was either correct or incorrect (off by 1 or 2), and participants had to indicate if the test sum was correct or not.

Metabolic data were collected using a metabolic cart using a mask attached to the face along with a heart rate monitor attached to the chest. The metabolic cart included gas analyzers in order to measure the concentrations of oxygen and carbon dioxide, and respiratory volume was measured using a respiratory pneumotachograph. The metabolic cart recorded variables every 5 sec as an average over the preceding 5 sec; therefore, for our analyses of the relationships involving our three candidates for the denominator of *NES*(*t*), we averaged all variables over all correct trials at each level of difficulty for each participant. Energy expended was calculated as discussed above.

EEG data were collected using a 64-channel system, with analog data digitized at a 1 KHz sampling rate. Data were preprocessed following a recently published pipeline by Delorme (2023). Variables were extracted for trials with correct responses and averaged at each level of difficulty for each participant to be consistent with what was allowable by the metabolic data. The values of GFP were calculated as noted above.^6^ The values for Σ(*α*) and Σ(*γ*) were calculated by performing a wavelet transform, positively rectifying the result, and summing power values across the two ranges of frequency.

### Relationships with metabolic energy expenditure

In order to assess the possible relationships of *GFP,* Σ(*α*), and Σ(*γ*) with metabolic energy expenditure, we first converted all variables to *Z*-scores at the individual subject level. We then averaged the *Z*-scores and examined the correlations between actual energy expended and each of the candidate indicators. Scatter plots for the relationships are presented in the three panels of Figure 4 and the calculated correlation coefficients are presented in Table 1. There were reliable relationships between energy expended and both *GFP* and Σ(*γ*) but no relationship with Σ(*α*). This is perhaps unsurprising as power in *γ* band is more frequently interpreted with respect to effortful attention than is power in *α* band. The correlations for *GFP* and Σ(*γ*) were almost identical for the two groups of participants.

**Figure 4.**
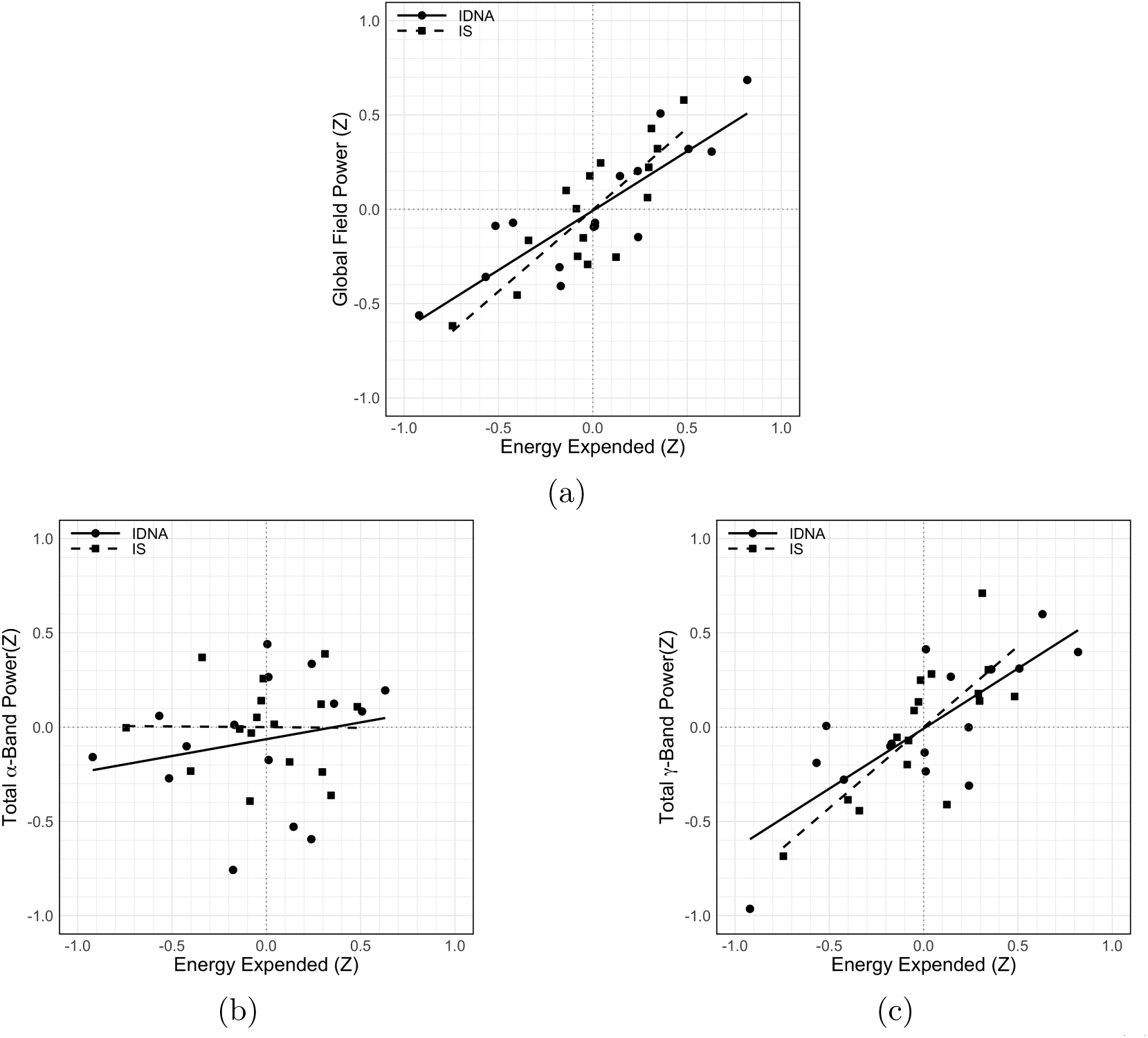
Scatter plots of the relationships between metabolic energy expended and (a) GFP, (b) Σ(α), and (c) Σ(γ). All variables are averaged Z-scores at each level of difficulty. Regression lines are plotted separately for the IS and IDNA participants.

**Table 1.** Correlations between metabolic energy expended and the three possible indicators, separately for the two groups of participants. Note: IDNA = iron deficient not anemic, IS = iron sufficient.

| Group | GFP(Z) | | $\Sigma(\alpha)(Z)$ | | $\Sigma(\gamma)(Z)$ | |
| --- | --- | --- | --- | --- | --- | --- |
| | $r$ | $p$ | $r$ | $p$ | $r$ | $p$ |
| IDNA | 0.86 | < 0.001 | 0.45 | 0.083 | 0.78 | < 0.001 |
| IS | 0.83 | < 0.001 | -0.01 | 0.971 | 0.77 | < 0.001 |

To determine whether the relationships between energy expended and *GFP* and Σ(*γ*) would be better explained by the variations in difficulty, we separately regressed *GFP* and Σ(*γ*) onto energy expended, set size, and number of segments using stepwise model selection to select the smallest number of predictors that accounted for the most variance. The results are presented in Table 2. Here it can be seen, that for these measures, the only significant predictor was metabolic energy expended. However, the proportion of variance accounted for by the relationship between energy expended and *GFP* was roughly twice that for the relationship between energy expended and Σ(*γ*).

**Table 2.** Results of regressing GFP (Z) and Σ(γ)(Z) onto the set of possible predictors using stepwise model selection. Note: EE = energy expended.

| Predicted<br>variable | Predictor | $\hat{\beta}$ | $SE$ | $p$ | $R^2$ |
| --- | --- | --- | --- | --- | --- |
| $GFP(Z)$ | $EE(Z)$ | 0.76 | 0.032 | $< 0.001$ | 0.553 |
| $\Sigma(\gamma)(Z)$ | $EE(Z)$ | 0.52 | 0.040 | $< 0.001$ | 0.264 |

At this point, we have demonstrated that *GFP* is superior to both Σ(*α*) and Σ(*γ*) in terms of the strength of its relationship to metabolic energy expenditure. And the strength of that relationship suggests the utility of using *GFP* —preferably *GFP* (*t*)—in constructing *NES*(*t*). However, we have yet to demonstrate that doing so provides unique insights into differences in processing as a function of some differences in state. That is the issue to which we turn next.

## Application of *NES*(*t*)

The possibility that the expenditure of brain energy in the service of perceptual and cognitive work is reduced in cases of iron deficiency, as is true for the expenditure of physical energy (Haas & Brownlie, 2001; R. Li et al., 1994; Luna et al., 2020; Y. I. Zhu & Haas, 1997; Y. Zhu & Haas, 1998) is suggested by the published data from the study just considered. Specifically, across the range of difficulty manipulations, women in the IDNA group were slower and less accurate than the women in the IS group. Yet, across the majority of this same range, women in the IDNA group were expending *the same* amount of energy as the women in the IS group. Essentially, their return on the same level of expenditure of energy was lower, suggesting that they were less efficient.

To determine if we could document a difference in efficiency as a function of iron status, we applied *NES*(*t*) to previously unpublished data from another previous study (Newbolds & Wenger, 2024). This was a study that examined the potential for using the electroretinogram (ERG) to assess differences between women who were IDNA and women who were IS that would indicate differences in central dopamine. The question was motivated by a set of findings in animal models (e.g., Ashkenazi, Ben-Shachar, & Youdim, 1982; Bianco, Wiesinger, Earley, Jones, & Beard, 2008; Erikson, Jones, Hess, Zhang, & Beard, 2001; Nelson, Erikson, Piñero, & Beard, 1997; Unger, Wiesinger, Hao, & Beard, 2009; Unger, Bianco, Jones, Allen, & Earley, 2014) which has shown that iron deficiency produces reductions in levels of dopamine and levels of dopamine transporters D1 and D2. The data from Newbolds and Wenger (2024) is drawn from a contrast detection task, performance on which has been shown to be affected by iron deficiency (Wenger, Murray-Kolb, et al., 2017) and by variations in central dopamine (Bodis-Wollner et al., 1987; Chen et al., 2003; Langheinrich et al., 2000).

### Description of the experimental methods

As with the study discussed in the previous section, two groups of women were recruited after being screened by way of blood assays: IDNA women and IS women, with the same inclusion criteria as in Wenger et al. (2017). The experimental task was based on a task and stimuli we have previously used in the study of visual perceptual learning (e.g., Wenger, Copeland, Bittner, & Thomas, 2008; Wenger & Rhoten, 2020). The stimuli were grayscale Gabor patches that were fixed in orientation and spatial frequency and that varied in (Michelson) contrast from 1-80% in steps of 0.1%. Trials were self-initiated by participants and began with the presentation of a fixation cross, followed by a pattern pre-mask, the test stimulus at the current level of contrast, and then a pattern post-mask. Participants had to respond as quickly as possible as to whether or not they perceived contrast to be present in the test stimulus. Contrast was adjusted according to a three-down/one-up staircase, with reductions occurring after three correct responses and increases happening after one incorrect response. Concurrent EEG and ERG data were collected during the performance of the task, using a 128-channel system with analog data digitized at 1 KHz.

Estimates of *h*(*t*) for each participant were obtained using RTs for correct responses using Kaplan-Meier estimates (e.g., Jager, Van Dijk, Zoccali, & Dekker, 2008) after being placed in 10 ms bins running from 200 to 1000 ms. The EEG data were preprocessed as above (Delorme, 2023) and *GFP* (*t*) was calculated as above but in this case in 10 ms bins running from 200 to 1000 ms after the onset of the stimulus. The *NES*(*t*) was then calculated for each participant as the ratio of *h*(*t*) to *GFP* (*t*), representing the relationship between the instantaneous measure of work accomplished and the instantaneous proxy measure of brain energy expended.^7^

### Differences in neural efficiency

To put the results in context, the women in the IDNA group had significantly higher contrast detection thresholds (15. 3%) than the women in the IS group (2.7%). This is consistent with findings from one of our field studies (Wenger, Murray-Kolb, et al., 2017) that, among other findings, demonstrated that repleting women with iron deficiency resulted in a significant decrease in their contrast detection thresholds.

The three panels of Figure 5 present the components of *NES*(*t*) and *NES*(*t*) as a function of iron status. Consider first the *h*(*t*) data in Figure 5a. Here it can be seen that the IS women were able to accomplish more work than were the IDNA women. However, the *GFP* (*t*) data (Figure 5b) show that the IS women accomplished more work while expending less energy than the IDNA women. Consequently, the *NES*(*t*) data (Figure 5c) confirm that IDNA women were less efficient in accomplishing their perceptual work than were the IS women. This difference in efficiency replicates a similar difference in efficiency as a function of iron status in the performance of a category learning task (Rhoten, Wenger, & De Stefano, 2024).

**Figure 5.**
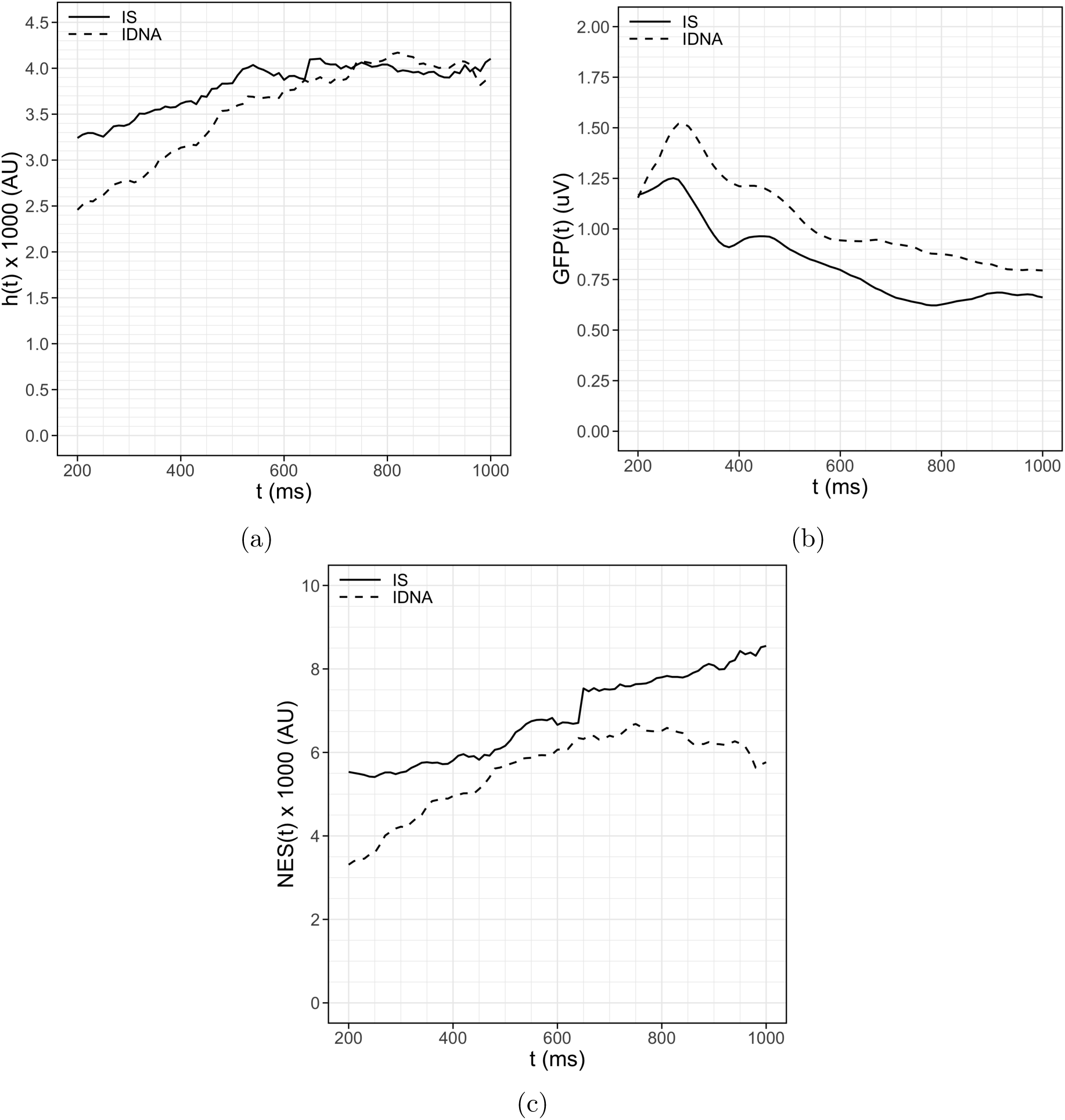
Neurocognitive efficiency in the contrast detection threshold data from Newbolds and Wenger (2024), as a function of iron status: (a) h(t), (b) GFP (t), (c) NES(t). Note: IDNA = iron deficient not anemic, IS = iron sufficient.

We next examined the extent to which *NES*(*t*) and its components were related to the iron biomarkers that were collected. Pairwise correlations between *h*(*t*), *GFP* (*t*), and *NES*(*t*) and a set of the biomarkers are presented in Table 3. Here it can be seen that *h*(*t*) was positively related to sFt, in terms of both raw values and log_10_ transformed values.^8^ This is the form of iron measured to index the level of iron stored in the body and is the form of iron that has been related to the synthesis and regulation of dopamine in the central nervous system (Beard & Connor, 2003; Erikson et al., 2001; Unger et al., 2007; Youdim & Yehuda, 2000). *GFP* (*t*) was negatively related to sFt (raw and transformed), and was also negatively related to mean corpuscular hemoglobin concentration (MCHC). The latter is critical with respect to interpreting *GFP* (*t*) in terms of energy expenditure, as MCHC is one of the iron biomarkers indicative of oxygen transport capacity. *NES*(*t*)—composed as it is of measures of work accomplished and energy expended—was positively related to both sFt and MCHC, suggesting that higher efficiency depends on both the aspect of iron that is related to the integrity of dopaminergic signaling (i.e., signal quality) and the aspect of iron that is related to the ability expend energy (i.e., by way of oxygen transport to and within the brain). In addition, *NES*(*t*) was negatively related to threshold, indicating that lower thresholds were related to higher neurocognitive efficiency.

**Table 3.** Pairwise correlations involving the NES and its components (h(t) and GFP), the blood iron biomarkers, and the contrast detection threshold. Cells shaded in gray indicate correlations that were reliably different from 0. Note: Hb = hemoglobin, sFt = serum ferritin, MCHC = mean corpuscular hemoglobin concentration, RDW = red blood cell distribution width.

|  | hazard |  | GFP |  | NES |  |
| --- | --- | --- | --- | --- | --- | --- |
|  | *r* | *p* | *r* | *p* | *r* | *p* |
| Hb (g/dL) | 0.26 | 0.109 | -0.12 | 0.488 | 0.16 | 0.359 |
| sFt (ng/mL) | 0.48 | 0.002 | -0.46 | 0.005 | 0.42 | 0.011 |
| $\log_{10}(\text{sFt})$ | 0.40 | 0.012 | -0.58 | < 0.001 | 0.45 | 0.006 |
| MCHC (g/dL) | 0.26 | 0.107 | -0.37 | 0.027 | 0.33 | 0.047 |
| RDW (%) | 0.01 | 0.955 | 0.33 | 0.058 | -0.20 | 0.253 |
| Threshold (%) | -0.21 | 0.204 | 0.26 | 0.131 | -0.66 | < 0.001 |

All of the results reported so far indicate that neurocognitive efficiency may mediate the relationship between iron status and perceptual performance, as measured in the contrast detection threshold task. To do this, we fit a set of alternative conditional process models (Hayes, 2018) to the data. The models considered were (a) a simple mediation model in which sFt was the predictor, threshold was the outcome, and *NES* was the mediator; variations on that simple model in which MCHC was (b) a covariate or (c) a moderator; and (d) a simple mediation model in which a composite of sFt and MCHC was the predictor, threshold was the outcome, and *NES* was the mediator. Since the conditional process models do not work with time series, we needed first to convert *NES*(*t*)*, h*(*t*) and *GFP* (*t*) to single values. We did this at the individual participant level by taking an interval of 100 ms around the participant’s median RT for correct responses and calculating the average value for each of the three variables within that interval. We then converted all variables to *Z*-scores. We formed a composite iron variable by taking the mean of the *Z*-scores for sFt and MCHC. The model that provided the best account of the data (in terms of largest values of *R*^2^) was the model in which the iron composite variable was the predictor. Should we find evidence that *NES* does mediate the relationship between iron status and detection threshold, we also need to know whether *NES* provides a better account of the overall relationship than does the hazard or *GFP* alone. Consequently, we also fit the four possible models using the hazard and *GFP* as the mediators.

For all three possible mediators, the form of the model that provided the best account of the data was the one in which the composite iron variable was the predictor. The results of fitting that model with the three candidate predictors are presented in Table 4. Of note first is the fact that the model in which *NES* was the mediator accounted for a far larger proportion of the variance (*R*^2^ = 0.44) than did the models in which either the hazard (0.08) or *GFP* (0.11) were the mediators. the second thing to note is that, when the complete indirect path was taken into account, the direct relationship between the iron composite variable and threshold was no longer significant. That is, the relationship between iron status and perceptual performance was completely mediated by neurocognitive efficiency.

**Table 4.**
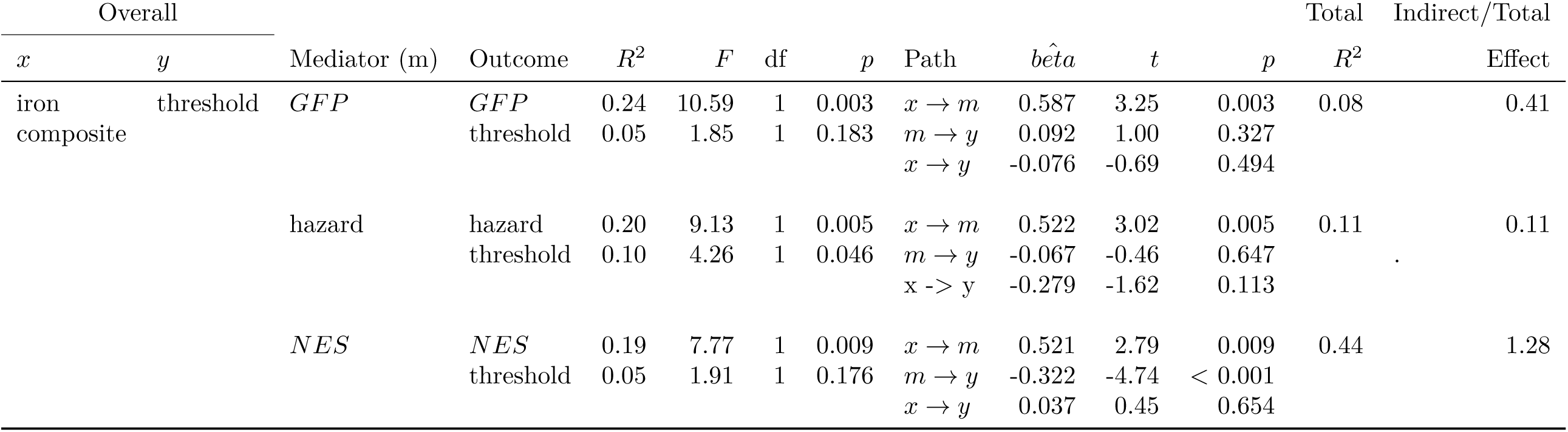
Parameters for the best-fitting mediation models involving NES and its components.

## Discussion

In order to perform the work supporting perception and cognition, the brain must expend energy and, just as is the case with physical work, the efficiency with which that energy is expended can vary. That efficiency can be adequately characterized by comparing the amount of work accomplished to the amount of energy expended. This set up the two-part problem that we sought to solve.

First, we needed to have a measure of perceptual and cognitive work accomplished. For this, we drew on a long line of work (beginning, e.g., with Townsend & Ashby, 1978) that has made fruitful use of the hazard function on the reaction time distribution, along with its integral, the former which can be interpreted as an instantaneous measure of the amount of work being done (the intensity of processing) and the latter which can be interpreted as a cumulative measure of amount accomplished. We noted how the integrated hazard function serves the critical role in the characterization of workload capacity in systems factorial technology, by allowing the comparison of cumulative work accomplished across levels of demand (Townsend & Nozawa, 1995; Townsend & Wenger, 2004).

Second, we needed to have a usable measure of brain energy expenditure during the performance of perceptual and cognitive work. This led us to consider the extent to which measures on EEG might be useful, as the time-course of this type of data is commensurate with the time-course of perceptual and cognitive processing. This led to consider one measure in the time domain—global field power (GFP)—and two measures in the time-frequency domain—summed power in the *α* and *γ* bands (Σ(*α*), Σ(*γ*). Having both a measure of work accomplished and potential measures of energy expended, we then could model our characterization of neurocognitive energetic efficiency on the characterization of physical energetic efficiency: as a ratio with the measure of work accomplished in the numerator and the possible measures of energy expended in the denominator.

This possibility of quantifying neurocognitive efficiency was critically dependent on being able to show that at least one of our possible EEG measures of energy expended had a reliable relationship with metabolic energy expended during the performance of a perceptual or cognitive task. It is rare to have measures of metabolic energy expended in perceptual and cognitive studies, but we were fortunate to be able to draw on unpublished data from a previous study (Wenger, DellaValle, et al., 2017) in which we collected concurrent behavioral, EEG, and metabolic data in a cognitive analog to a physical exercise test. Using these data, we were able to show that both *GFP* and Σ(*γ*) were reliably related to metabolic energy, with the strongest relationship being with *GFP*

This result determined the form of *NES*(*t*) that we applied to unpublished data from another recent study (Newbolds & Wenger, 2024) comparing the performance of women of reproductive age who were iron sufficient to those who were iron deficient but not anemic. We demonstrated that differences in iron status were reliably related to differences in neurocognitive efficiency, that neurocognitive efficiency was reliably related to measures of iron that themselves are related to both the integrity of neurotransmission (signal quality) and oxygen transport (energy expenditure), and that neurocognitive efficiency completely mediated the relationship between iron status and perceptual performance.

With respect to strengths and avenues for future work, we see the potential for broad applicability of this new measure. With respect to theory, an obvious and necessary step will be to formally connect this measure to our characterizations of capacity, including those specified both in terms of RTs alone and performance conditioned on accuracy (e.g. Altieri, Townsend, & Wenger, 2014; Townsend & Altieri, 2012; Townsend & Eidels, 2011; Townsend & Nozawa, 1995; Townsend & Wenger, 2004). In terms of basic science, having a measure of neuocognitive efficiency along with indirect measures of dopamine opens the possibility of quantifying two factors that may contribute to a mechanistic understanding of how iron deficiency extracts costs in perceptual and cognitive performance. In terms of more applied questions, applications that might be of interest would include comparison of novices and experts, measurement of change as a function of skill learning, measurement of normal and disease-related changes as a function of age, and the effects of physical interventions such as exercise on cognitive performance (as in Fu & Yang, 2024).

There are a few caveats that need to be considered. The first and most obvious is that the finding of a positive relationship between metabolic energy expended and *GFP* needs to be replicated. Of concern here is the fact that we used large variations in workload—much larger than those used in earlier studies of the effect of cognitive workload on metabolic measures (Garde et al., 2002; Petrek, 2008, 2009). It may be that the positive relationship that we obtained is only observable at high levels of demand. The second caveat is that all of our results were obtained in comparisons between ID and IDNA women, suggesting a potential limit on generalizability to other populations, ages, and biological sex. The third caveat is that there may be other measures in the time frequency, such as the *θ/β* ratio (e.g, Angelidis, Hagenaars, van Son, van der Does, & Putman, 2018) that need to be assessed. However, should these questions be addressed and positive outcomes obtained, the *NES*(*t*) would then have the potential to be used in contexts in which the energetic demands of the experimental tasks have been given either little or no attention and for experimentalists and theoreticians to work to ground characterizations of perceptual and cognitive work on the physical expenditure of energy (i.e., the thermodynamics of perception and cognition, Kringelbach, Perl, & Deco, 2024).

## Footnotes

1 We thank Joe Lappin for pointing this out.

2 Though intended to be applied to human information processing, these four characteristics can be considered for *any* information processing systems, including those of other species and engineered systems.

3 Biofortification is the process of selectively breeding plants to increase a specific nutrient, iron in this case, and can be distinguished from genetic engineering.

4 A common misconception is that iron deficiency and anemia are the same thing, when in fact they can be thought of as orthogonal, in that it is possible to be anemic without being iron deficient and vice versa.

5 We opt not to consider measures in the frequency domain as we wish to retain temporal information.

6 Note that as we are averaging at each level of difficulty it is not appropriate to refer to any of our candidate measures as a function of time.

7 We acknowledge that we are using the term “instantaneous” somewhat loosely, as the data in fact were separated into 10 ms bins.

8 Values of sFt are not always normally distributed,so it is standard practice to present both the raw and the transformed values.

## References

Altieri, N., Townsend, J. T., & Wenger, M. J. (2014). A measure for assessing the effects of audiovisual speech integration. Behavior Research Methods, 46 (2), 406–415.

Ames, A. (2000). CNS energy metabolism as related to function. Brain Research Reviews, 34 (1-2), 42–68.

Anemia. (2024). Retrieved from https://www.who.int/health-topics/anaemia (Accessed on 25 May, 2024)

Angelidis, A., Hagenaars, M., van Son, D., van der Does, W., & Putman, P. (2018). Do not look away! Spontaneous frontal EEG theta/beta ratio as a marker for cognitive control over attention to mild and high threat. Biological Psychology, 135, 8–17.

Ashkenazi, R., Ben-Shachar, D., & Youdim, M. B. (1982). Nutritional iron and dopamine binding sites in the rat brain. Pharmacology Biochemistry and Behavior, 17, 43–47.

Beard, J. L., & Connor, J. R. (2003). Iron status and neural functioning. Annual Review of Nutrition, 23 (1), 41–58.

Berka, C., Levendowski, D. J., Lumicao, M. N., Yau, A., Davis, G., Zivkovic, V. T., … Craven, P. L. (2007). Correlates of task engagement and mental workload in vigilance, learning, and memory tasks. *Aviation*, Space, and Environmental Medicine, 78 (5), B231–B244.

Bianco, L. E., Wiesinger, J., Earley, C. J., Jones, B. C., & Beard, J. L. (2008). Iron deficiency alters dopamine uptake and response to l-dopa injection in sprague–dawley rats. Journal of Neurochemistry, 106 (1), 205–215.

Blaha, L. M., Busey, T. A., & Townsend, J. T. (2009). An LDA approach to the neural correlates of configural learning. In Proceedings of the 31st annual conference of the cognitive science society. Austin, TX.

Blaha, L. M., & Houpt, J. W. (2015). An extension of workload capacity space for systems with more than two channels. Journal of Mathematical Psychology, 66, 1–5.

Bodis-Wollner, I., Marx, M. S., Mitra, S., Bobak, P., Mylin, L., & Yahr, M. (1987). Visual dysfunction in parkinson’s disease: Loss in spatiotemporal contrast sensitivity. Brain, 110 (6), 1675–1698.

Borghetti, L., Morris, M. B., Jack Rhodes, L., Haubert, A. R., & Veksler, B. Z. (2021). Gamma oscillations index sustained attention in a brief vigilance task. In Proceedings of the human factors and ergonomics society annual meeting (Vol. 65, pp. 546–550).

Centanni, T. M., Halpern, A. R., Seisler, A. R., & Wenger, M. (2020). Context-dependent neural responses to minor notes in frontal and temporal regions distinguish musicians from nonmusicians. *Cognitive, Affective*, & Behavioral Neuroscience, 20, 551–564.

Chechile, R. A. (2003). Mathematical tools for hazard function analysis. Journal of Mathematical Psychology, 47 (5-6), 478–494.

Chechile, R. A. (2011). Properties of reverse hazard functions. Journal of Mathematical Psychology, 55 (3), 203–222.

Chechile, R. A. (2022). Using hazard and surrogate functions for understanding memory and forgetting. Applied Math, 2 (4), 518–546.

Chen, Y., Levy, D. L., Sheremata, S., Nakayama, K., Matthysse, S., & Holzman, P. S. (2003). Effects of typical, atypical, and no antipsychotic drugs on visual contrast detection in schizophrenia. American Journal of Psychiatry, 160 (10), 1795–1801.

Cooper, C. B., & Storer, T. W. (2001). Exercise testing and interpretation: a practical approach. Cambridge University Press.

Cowan, N. (2012). Working memory capacity. Psychology Press.

Cox, D. R. (1962). Renewal theory. London: Methuen.

Cunningham, C. A., Yassa, M. A., & Egeth, H. E. (2015). Massive memory revisited: Limitations on storage capacity for object details in visual long-term memory. Learning & Memory, 22 (11), 563–566.

Dasari, D., Shou, G., & Ding, L. (2017). ICA-derived EEG correlates to mental fatigue, effort, and workload in a realistically simulated air traffic control task. Frontiers in Neuroscience, 11, 297.

DellaValle, D. M., & Haas, J. D. (2014). Iron supplementation improves energetic efficiency in iron-depleted female rowers. Medicine and Science in Sports and Exercise, 46 (6), 1204–1215.

Delorme, A. (2023). EEG is better left alone. Scientific Reports, 13 (1), 2372.

de V. Weir, J. B. (1949). New methods for calculating metabolic rate with special reference to protein metabolism. Journal of Physiology, 109, 1–9.

Doesburg, S. M., Roggeveen, A. B., Kitajo, K., & Ward, L. M. (2008). Large-scale gamma-band phase synchronization and selective attention. Cerebral cortex, 18 (2), 386–396.

Erikson, K. M., Jones, B. C., Hess, E. J., Zhang, Q., & Beard, J. L. (2001). Iron deficiency decreases dopamine d1 and d2 receptors in rat brain. Pharmacology Biochemistry and Behavior, 69 (3), 409–418.

Fairclough, S. H., & Houston, K. (2004). A metabolic measure of mental effort. Biological Psychology, 66 (2), 177–90.

Finkelstein, M. (2008). Failure rate modelling for reliability and risk. Springer Science & Business Media.

Fisher, D. L. (1982). Limited-channel models of automatic detection: Capacity in scanning in visual search. Psychological Review, 89, 662–692.

Fisher, D. L. (1984). Central capacity limits in consistent mapping, visual search tasks: Four channels or more? Cognitive Psychology, 16, 449–484.

Fox, P. T., & Raichle, M. E. (1986). Focal physiological uncoupling of cerebral blood flow and oxidative metabolism during somatosensory stimulation in human subjects. Proceedings of the National Academy of Science USA, 83 (4), 1140–1144.

Fox, P. T., Raichle, M. E., Mintun, M. A., & Dence, C. (1988). Nonoxidative glucose consumption during focal physiologic neural activity. Science, 241 (4864), 462–464.

Fu, H.-L., & Yang, C.-T. (2024). Exploring the influence of a 4-week aerobic exercise intervention on cognitive control processes in young adults: An SFT and DDM study. Progress in Brain Research, 283, 193–229.

Fukuda, K., & Vogel, E. K. (2019). Visual short-term memory capacity predicts the “bandwidth” of visual long-term memory encoding. Memory & cognition, 47, 1481–1497.

Garde, A. H., Laursen, B., Jørgensen, A. H., & Jensen, B. R. (2002). Effects of mental and physical demands on heart rate variability during computer work. European Journal of Applied Physiology, 87, 456–61.

Haas, J. D. (2006). The effects of iron deficiency on physical performance. Food and Nutrition Board, Institute of Medicine. The National Academies Press, Washington, DC, 451–461.

Haas, J. D., & Brownlie, T. (2001). Iron deficiency and reduced work capacity: A critical review of the research to determine a causal relationship. Journal of Nutrition, 131, 676S–688S.

Hall, C. N., Klein-Flügge, M. C., Howarth, C., & Attwell, D. (2012). Oxidative phosphorylation, not glycolysis, powers presynaptic and postsynaptic mechanisms underlying brain information processing. Journal of Neuroscience, 32 (26), 8940–8951.

Halpern, A., Martin, J., & Reed, T. (2008). An ERP study of major-minor classification in melodies. Music Perception, 25 (3), 181–191.

Hayes, A. F. (2018). Introduction to mediation, moderation, and conditional process analysis: A Regression-based approach (third ed.). New York: Guilford Press.

Heathcote, A., Coleman, J. R., Eidels, A., Watson, J. M., Houpt, J., & Strayer, D. L. (2015). Working memory’s workload capacity. Memory & cognition, 1–17.

Houpt, J. W., & Townsend, J. T. (2012). Statistical measures for workload capacity analysis. Journal of Mathematical Psychology, 56 (5), 341–355.

Hyder, F. (2004). Deriving changes in cmr*_O_*_2_ from calibrated fMRI. In R. G. Shulman & D. L. Rothman (Eds.), Brain energetics and neuronal activity: Applications to fMRI and medicine (p. 147–171). West Sussex, England: Wiley.

Hyder, F., & Blumenfeld, H. (2004). Relationship between cmr*_O_*_2_ and neuronal activity. In Brain energetics and neuronal activity: Applications to fMRI and medicine (p. 173–194). West Sussex, England: Wiley.

Hyder, F., Rothman, D. L., & Shulman, R. G. (2002). Total neuroenergetics support localized brain activity: Implications for the interpretation of fMRI. Proceedings of the National Academy of Science USA, 99 (16), 10771–10776.

Jager, K. J., Van Dijk, P. C., Zoccali, C., & Dekker, F. W. (2008). The analysis of survival data: the Kaplan–Meier method. Kidney International, 74 (5), 560–565.

Jung, T.-P., & Lee, T.-W. (2007). Applications of independent component analyses to electroencephelography. In M. J. Wenger & C. Schuster (Eds.), Statistical and process models for cognitive neuroscience and aging (p. 313–334). Mahwah, NJ: Erlbaum.

Kail, R., & Salthouse, T. A. (1994). Processing speed as mental capacity. Acta Psychologica, 86, 199–225.

Kringelbach, M. L., Perl, Y. S., & Deco, G. (2024). The thermodynamics of mind. Trends in Cognitive Sciences.

Landau, A. N., Esterman, M., Robertson, L. C., Bentin, S., & Prinzmetal, W. (2007). Different effects of voluntary and involuntary attention on EEG activity in the gamma band. Journal of Neuroscience, 27 (44), 11986–11990.

Langheinrich, T., van Elset, L. T., Lagreze, W. A., Bach, M., Lucking, C. H., & Greenlee, M. W. (2000). Visual contrast response functions in Parkinson’s disease: evidence from electroretinograms, visually evoked potentials and psychophysics. Clinical Neurophysiology, 111, 66–74.

Lappin, J. S., & Bell, H. H. (2021). Form and function in information for visual perception. i-Perception, 12 (6), 20416695211053352.

Lappin, J. S., Morse, D. L., & Seiffert, A. E. (2016). The channel capacity of visual awareness divided among multiple moving objects. *Attention, Perception*, & Psychophysics, 78, 2469–2493.

Lappin, J. S., Seiffert, A. E., & Bell, H. H. (2020). A limiting channel capacity of visual perception: Spreading attention divides the rates of perceptual processes. *Attention, Perception*, & Psychophysics, 82, 2652–2672.

Laughlin, S. B., & Attwell, D. (2004). Neural energy consumption and the representation of mental events. In R. G. Shulman & D. L. Rothman (Eds.), Brain energetics and neuronal activity: Applications to fMRI and medicine (p. 111–124). West Sussex, England: Wiley.

Lehmann, D., & Skrandies, W. (1980). Reference-free identification of components of checkerboard-evoked multichannel potential fields. Electroencephalography and Clinical Neurophysiology, 48 (6), 609–621.

Lin, A. L., Fox, P. T., Hardies, J., Duong, T. Q., & Gao, J. H. (2010). Nonlinear coupling between cerebral blood flow, oxygen consumption, and atp production in human visual cortex. Proceedings of the National Academy of Sciences, 107, 8446–8451.

Little, D., Altieri, N., Yang, C., & Fific, M. (2017). Systems factorial technology: A theory driven methodology for the identification of perceptual and cognitive mechanisms. Elsevier Science & Technology Books. Retrieved from https://books.google.com/books?id=OQICMQAACAAJ

Longo, L., Wickens, C. D., Hancock, G., & Hancock, P. A. (2022). Human mental workload: A Survey and a novel inclusive definition. Frontiers in Psychology, 13, 883321.

Luna, S. V., Pompano, L. M., Lung’aho, M., Gahutu, J. B., & Haas, J. D. (2020). Increased iron status during a feeding trial of iron-biofortified beans increases physical work efficiency in rwandan women. The Journal of Nutrition, 150 (5), 1093–1099.

McGill, W. J., & Gibbon, J. (1965). The general gamma distribution and reaction times. Journal of Mathematical Psychology, 2, 1–18.

Mizokuchi, K., Tanaka, T., Sato, T. G., & Shiraki, Y. (2024). Alpha band modulation caused by selective attention to music enables eeg classification. Cognitive Neurodynamics, 18 (3), 1005–1020.

Nelson, C., Erikson, K., Piñero, D. J., & Beard, J. L. (1997). In vivo dopamine metabolism is altered in iron-deficient anemic rats. The Journal of Nutrition, 127 (12), 2282–2288.

Newbolds, S. F., & Wenger, M. J. (2024). Assessing the pattern electroretinogram as a proxy measure for dopamine in the context of iron deficiency. Nutritional Neuroscience, 1–12. doi: 10.1080/1028415X.2024.2304943

Noah, S., Powell, T., Khodayari, N., Olivan, D., Ding, M., & Mangun, G. R. (2020). Neural mechanisms of attentional control for objects: decoding EEG alpha when anticipating faces, scenes, and tools. Journal of Neuroscience, 40 (25), 4913–4924.

Nunez, P. L., & Srinivasan, R. (2006). Electric fields of the brain: The neurophysics of EEG (second ed.). Oxford: Oxford.

Oberauer, K., Farrell, S., Jarrold, C., & Lewandowsky, S. (2016). What limits working memory capacity? Psychological Bulletin, 142 (7), 758.

Özugur, S., Kunz, L., & Straka, H. (2020). Relationship between oxygen consumption and neuronal activity in a defined neural circuit. BMC Biology, 18, 1–16.

Petrek, J. (2008). Pictorial cognitive task resolution and dynamics of event-related potentials. Biomed Pap Med Fac Univ Palacky Olomouc Czech Repub, 152, 223–230.

Petrek, J. (2009). Pictorial cognitive task resolution and expired minute ventilation, oxygen consumption, carbon dioxide production and heart rate. Biomed Pap Med Fac Univ Palacky Olomouc Czech Repub, 153, 131–136.

Raichle, M. E., & Gusnard, D. A. (2002). Appraising the brain’s energy budget. Proceedings of the National Academy of Science USA, 99 (16), 10237–10239.

Rhoten, S. E., Wenger, M. J., & De Stefano, L. A. (2024). Iron deficiency negatively affects behavioral measures of learning, indirect neural measures of dopamine, and neural efficiency. Cognitive, Affective, and Behavioral Neuroscience. (in press)

R. Li, R., Chen, X., Yan, H., Deurenberg, P., Garby, L., & Hautvast, J. G. A. H. (1994). Functional consequences of iron supplementation in iron-deficient female cotton workers in Beijing, China. American Journal of Clinical Nutrition, 59, 908–913.

Shannon, C. E. (1948). A mathematical theory of communication. The Bell System Technical Journal, 27, 379–423.

Shannon, C. E., & Weaver, W. (1963). The mathematical theory of communication. Urbana, IL: University of Illinois Press.

Skrandies, W. (1989). Data reduction of multichannel fields: Global field power and principal components analysis. Brain Topography, 2, 73–80.

Skrandies, W. (1990). Global field power and topographic similarity. Brain Topography, 3, 137–141.

Skrandies, W., & Lehmann, D. (1982). Spatial principal components of multichannel maps evoked by lateral visual half-field stimuli. Electroencephalography and Clinical Neurophysiology, 54 (6), 662–667.

Sokoloff, L. (2004). Energy metabolism in neural tissues in vivo at rest and in functionally altered states. In R. G. Shulman & D. L. Rothman (Eds.), Brain energetics and neuronal activity: Applications to fMRI and medicine (p. 11–30). West Sussex, England: Wiley.

Souza, R. H. C. e., & Naves, E. L. M. (2021). Attention detection in virtual environments using EEG signals: a scoping review. Frontiers in Physiology, 12, 727840.

Sternberg, S. (1966). High-speed scanning in human memory. Science, 153 (3736), 652–654.

Townsend, J. T., & Altieri, N. (2012). An accuracy-response time capacity assessment function that measures performance against standard parallel predictions. Psychological Review, 119 (3), 500–516.

Townsend, J. T., & Ashby, F. G. (1978). Methods of modeling capacity in simple processing systems. In J. Castellan & F. Restle (Eds.), Cognitive theory (vol. 3) (p. 200–239). Hillsdale, NJ: Erlbaum.

Townsend, J. T., & Ashby, F. G. (1983). Stochastic modeling of elementary psychological processes. Cambridge: Cambridge University press.

Townsend, J. T., & Eidels, A. (2011). Workload capacity spaces: A unified methodology for response time measures of efficiency as workload is varied. Psychonomic Bulletin & Review, 659–681.

Townsend, J. T., & Nozawa, G. (1995). On the spatio-temporal properties of elementary perception: An investigation of parallel, serial, and coactive theories. Journal of Mathematical Psychology, 39, 321–359.

Townsend, J. T., & Wenger, M. J. (2004). A theory of interactive parallel processing: new capacity measures and predictions for a response time inequality series. Psychological Review, 111 (4), 1003.

Unger, E. L., Bianco, L. E., Jones, B. C., Allen, R. P., & Earley, C. J. (2014). Low brain iron effects and reversibility on striatal dopamine dynamics. Experimental Neurology, 261, 462–468.

Unger, E. L., Paul, T., Murray-Kolb, L. E., Felt, B., Jones, B. C., & Beard, J. L. (2007). Early iron deficiency alters sensorimotor development and brain monoamines in rats. The Journal of Nutrition, 137 (1), 118–124.

Unger, E. L., Wiesinger, J. A., Hao, L., & Beard, J. L. (2009). Dopamine D2 receptor expression is altered by changes in cellular iron levels in PC12 cells and rat brain tissue. The Journal of Nutrition, 138 (12), 2487–2494.

Von Der Heide, R. J., Wenger, M. J., Bittner, J. L., & Fitousi, D. (2018). Converging operations and the role of perceptual and decisional influences on the perception of faces: Neural and behavioral evidence. Brain and Cognition, 122, 59–75.

Wenger, M. J., Copeland, A. M., Bittner, J. L., & Thomas, R. D. (2008). Evidence for criterion shifts in visual perceptual learning: Data and implications. Perception & Psychophysics, 70, 1248–1273.

Wenger, M. J., DellaValle, D. M., Murray-Kolb, L. E., & Haas, J. D. (2017). Effect of iron deficiency on simultaneous measures of behavior, brain activity, and energy expenditure in the performance of a cognitive task. Nutritional Neuroscience, 22 (3), 196–206.

Wenger, M. J., Murray-Kolb, L. E., Nevins, J. E. H., Venkatramanan, S., Reinhart, G. A., Wesley, A., & Haas, J. D. (2017). Consumption of a double-fortified salt affects perceptual, attentional, and mnemonic functioning in women in a randomized controlled trial in india. The Journal of Nutrition, 147 (12), 2297–2308.

Wenger, M. J., & Rhoten, S. E. (2020). Perceptual learning produces perceptual objects. *Journal of Experimental Psychology: Learning*, Memory, and Cognition, 46 (*3*), 455–475.

Wickens, C. D. (1984). Processing resources in attention. In R. Parasuraman & R. Davies (Eds.), Varieties of attention (p. ??). New York: Academic.

Woodroofe, M. (1985). Estimating a distribution function with truncated data. The Annals of Statistics, 13 (1), 163–177.

Youdim, M. B., & Yehuda, S. (2000). The neurochemical basis of cognitive deficits induced by brain iron deficiency: Involvement of dopamine-opiate system. Cellular and Molecular Biology, 46 (3), 491–500.

Yu, J.-C., Chang, T.-Y., & Yang, C.-T. (2014). Individual differences in working memory capacity and workload capacity. Frontiers in Psychology, 5.

Zhu, Y., & Haas, J. D. (1998). Altered metabolic response of iron-depleted non-anemic women during a 15 km time trial. Journal of Applied Physiology, 84, 1768–1775.

Zhu, Y. I., & Haas, J. D. (1997). Iron depletion without anemia and physical performance in young women. American Journal of Clinical Nutrition, 66 (2), 334–41.

